# Fertility Gene Introns Harbor Transposable Elements that Shape Y-Loop Architecture

**DOI:** 10.64898/2026.07.08.737335

**Authors:** E.K. Beard, Jeffrey P Gamer, Amelie Raz, Mayu Inaba

## Abstract

Transposable elements (TEs) are powerful drivers of genome evolution, yet how they persist under selection and become incorporated into host regulatory networks remains poorly understood. In the *Drosophila* male germline, TEs are highly expressed during the spermatocyte stage, coinciding with activation of giant fertility genes on the Y chromosome. These genes contain megabase-scale introns enriched for repetitive DNA, and three of these genes form prominent nuclear structures known as Y-loops, providing a unique system to investigate gene regulation. Here, we show that multiple TEs expressed in spermatocytes are transcribed from the introns of Y-linked fertility genes. RNA fluorescence in situ hybridization (FISH) targeting several TEs, including *accord2*, *Juan*, and *HMS Beagle*, illuminates distinct nuclear regions corresponding to *kl-2*, *kl-3*, and *kl-5*, respectively. Genetic perturbation of these fertility gene loci or disruption of RNA-processing factors eliminates these TE transcripts, demonstrating that these TE sequences are embedded within Y chromosome-associated nascent transcripts rather than being independently transcribed. The identity and expression patterns of Y-loop-associated TEs vary extensively among closely related *Drosophila* species, consistent with the previously documented rapid evolution of Y-linked loci and suggesting that TEs may contribute to the genetic diversification of these giant fertility genes. We propose that continual turnover of repetitive elements within Y-linked introns provides a mechanism by which rapidly evolving repetitive DNA influences germline gene regulation, male fertility, and speciation.

## Introduction

Transposable elements (TEs) are a major source of genetic variation and genome innovation. By inserting into new genomic locations, altering gene structure, and reshaping regulatory networks, TEs continuously generate genomic changes that can serve as raw material for evolution. However, fundamental questions remain unresolved. How do TEs contribute to genome evolution and how do they persist under strong selective pressure? The answers likely lie in their tightly regulated activity during key developmental windows, especially in germ cells, where heritable genetic changes can occur.

A common feature across many organisms is that TEs are transiently reactivated during periods of epigenetic reprogramming, such as early embryogenesis, germline specification, and gametogenesis, when genome-wide chromatin remodeling temporarily relaxes repression ^1^. This reactivation may occur passively as a consequence of lost silencing, but some evidence suggests that certain TEs actively exploit these developmental windows by evolving regulatory features that synchronize their mobilization with host developmental transitions^2^.

In *Drosophila* males, robust TE expression is observed specifically during the spermatocyte stage of spermatogenesis, a period coinciding with the massive transcriptional activation of fertility genes ^3–5^. Several studies further indicate that at least a subset of these TEs remains mobile during spermatogenesis, implicating the male germline as a hotspot for TE propagation and genome diversification ^5–7^. However, the biological significance and mechanism for this stage-specific TE activation remain unclear. Is it merely a byproduct of the open chromatin environment required for the expression of fertility genes, or are TEs actively integrated into the regulatory circuitry governing spermatocyte development? Furthermore, what mechanisms balance their activity with genome integrity during germ cell differentiation?

Among the genes expressed in spermatocytes, the Y chromosome fertility genes are indispensable for male fertility and are of particular interest because several of them form massive lampbrush-like chromatin loops, known as Y-loops, that represent some of the largest actively transcribed gene structures in eukaryotic genomes.^8,9^. These genes contain extraordinarily large introns spanning megabases and are composed largely of simple repetitive sequences^10–12^. Consequently, their transcription and processing present exceptional challenges to the cellular gene-expression machinery ^13–15^. Successful expression of these loci requires coordinated regulation over long genomic distances and specialized mechanisms to ensure proper splicing and processing. Their dependence on such complex processing pathways likely makes them inherently sensitive to genetic and regulatory perturbations^15^. Consistent with this idea, both cis-regulatory features and trans-acting factors involved in Y-linked gene expression show rapid evolutionary divergence and can contribute to reproductive incompatibilities between species ^16–18^.

The intrinsic vulnerability of these genes may also make them particularly susceptible to the effects of TE insertions. Insertions within their already massive introns or changes in TE insertion/transcription that may influence chromatin organization and RNA processing could have disproportionate consequences for gene expression and male fertility. Because these genes are indispensable for reproduction, understanding how TEs interact with Y-linked fertility loci may provide fundamental insights into the mechanisms that drive genome evolution in natural populations.

Here, we show that a large proportion of TE transcripts are likely expressed within the introns of genes expressed in spermatocytes and are eliminated upon knocking down of a splicing factor *U2af38*. In particular, we show three that TEs, *Juan, HMS Beagle* and *accord2*, are expressed from introns of Y-linked fertility genes, *kl-3, kl-5*, and *kl-2* respectively. Moreover, we find that the expression or genome location of Y-loop-associated TEs vary significantly among closely related *Drosophila* species. These findings show that TE turnover within giant introns may serve as a mechanism through which rapidly evolving repetitive DNA shapes germline development, reproductive fitness, and ultimately, species divergence.

## Results

### Retrotransposon transcripts show distinct localization patterns in primary spermatocyte nuclei

We used RNA fluorescence in situ hybridization (FISH) to examine the expression of retrotransposons throughout the stages of spermatogenesis in the *Drosophila* testes (Figure 1A). We tested five TEs—*Juan*, *HMS Beagle*, *accord2*, *McClintock,* and *mdg4*—all of which have previously been shown to be expressed at high levels in SCs by RNA sequencing ^5^. Consistent with previous reports on the expression of TEs in the male germline ^4–6,19–21^, we found two principles which applied to all five TEs we tested. First, we primarily detect abundant transcripts in primary SCs, but detect limited amounts in early germ cells, spermatids, or somatic cells (Figure 1B-F). Second, TE transcripts are generally retained in the nucleus, with limited, if any, cytoplasmic signal (Figure 1G-P). Within the nucleus, however, each TE has its own distinct pattern of localization (Figure 1G-U).

**Figure 1.**
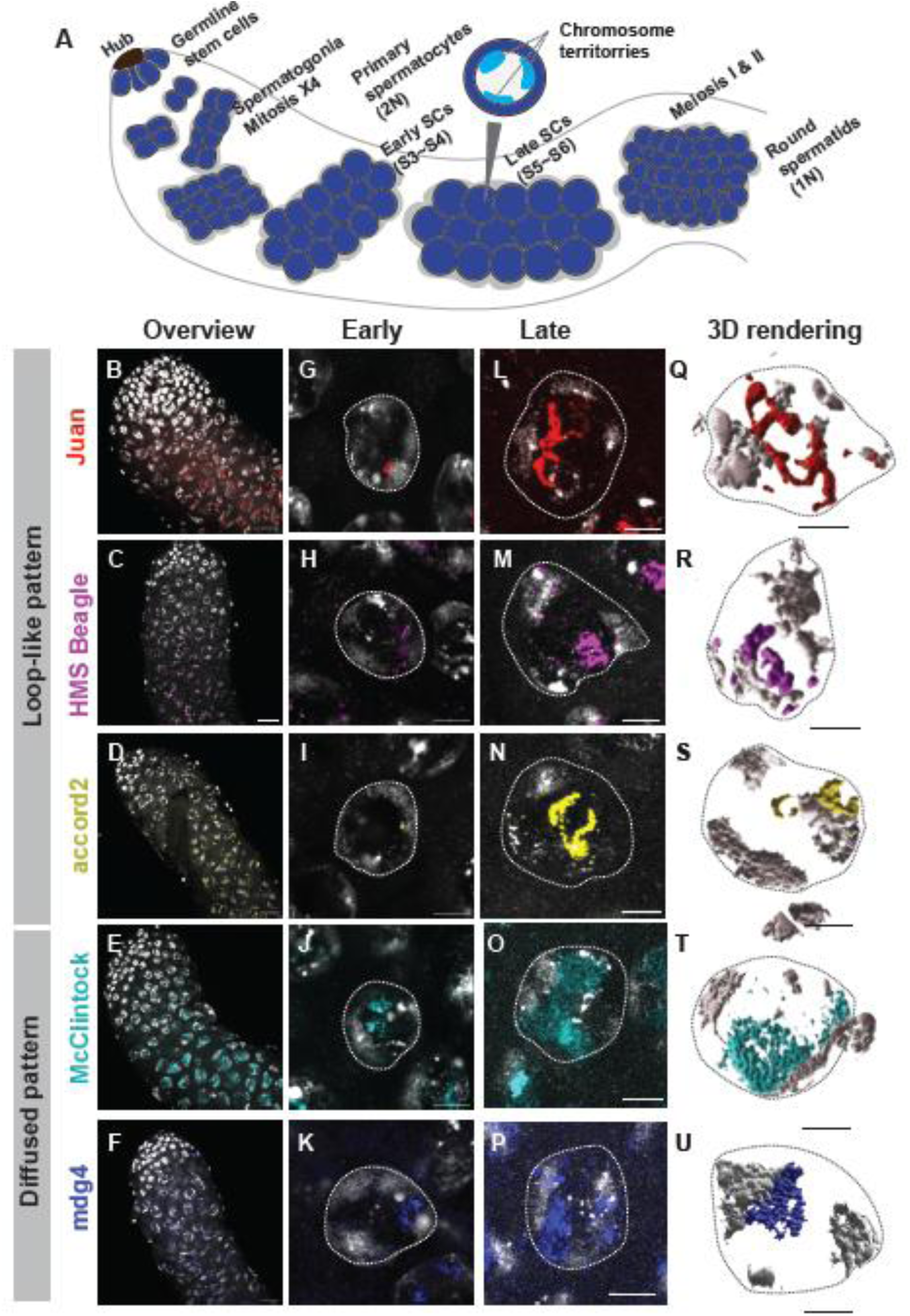
Retrotransposon transcripts show distinct localization patterns in primary spermatocyte nuclei. (**A**) The diagram shows the anatomy of the *Drosophila* testis and stage progression of spermatogenesis. (**B–F**) Representative RNA FISH images of testis tips using probes against the indicated TEs. Scale bars: 10 µm. (**G–K**) Representative images of early SCs (S3–S4) using probes against the indicated TEs. Scale bars, 5 µm. (**L–P**) Representative images of late SCs (S5–S6) using probes against the indicated TEs. Scale bars: 5 µm. (**Q-U**) 3D rendering of the images shown in **L-P** using Imaris software. White dotted lines encircle SC nuclei. *w1118* genotype was used for all images. DAPI (gray).

*Juan* is a non-LTR retrotransposon which is known to be recently active in *Drosophila melanogaster* ^22^. Staging of *Drosophila* SCs has been established by previous studies ^23^. *Juan* RNA is visible as a single punctum in early SCs (S3∼S4) (Figure 1C), closely associated with chromatin. In late SCs (S5∼S6), Juan transcript has increased and is organized in a string- or loop-like pattern throughout the nucleoplasm (Figure 1L, Q). Juan remains in this configuration throughout late SCs and suddenly disappears in M1. RNA from the LTR retrotransposon *HMS Beagle* follows a similar trajectory to *Juan*. The signal is small and closely associated with chromatin in early SCs (Figure 1H), and in late SCs, it accumulates in a loop-like pattern in the nucleoplasm, though in a smaller, less central area than Juan (Figure 1M, R). *accord2* is a LTR retrotransposon. Similarly, *accord2* RNA begins as a small punctum closely associated with chromatin in early SCs (Figure 1I). In late SCs, typically, two small loop-like accord2 signals are observed (Figure 1I, N).

The localization patterns of *McClintock (quasimodo2)* and *mdg4* (see ^24^), both LTR retrotransposons, differ markedly from those of the other transposable elements examined. In contrast to the characteristic loop-like pattern, RNA FISH signals for these elements appear within a small portion near the chromosomes in early spermatocytes (Figure 1J, K). This signal diffuses further in late spermatocytes, occupying a larger nuclear region (Figure 1O, T, P, U).

In summary, TEs are expressed in SC nuclei, and localization of TE transcript is element specific.

### TE transcripts originate from the Y chromosome

While TE transcription in SCs has been well established ^5^, it remains unclear whether TEs are transcribed independently or as part of host gene transcripts. Because nuclear RNA compartments are often determined by the genomic site of transcription^25^, the distinct nuclear localization patterns of TE transcripts led us to ask whether these patterns reflect their genomic origins. However, the multicopy nature of transposable elements and the challenges of assembling repetitive genomic regions make it difficult to assign TE expression to specific loci using bioinformatic approaches alone ^17^. Therefore, we combined genetic deletions and deficiency lines with RNA FISH to identify the genomic origins of TE transcription.

The Y chromosome contains genes with giant introns, including the fertility factors *kl-5, kl-3, kl-2*, *ORY (ks-1),* and *CCY (ks-2)* ^26^. An important feature of SC nuclear architecture is the formation of the Y-loops, lampbrush loop-like structures, originating from three of the Y chromosome gigantic genes. These three fertility factors, *kl-5, kl-3,* and *ORY* (*ks-1*) form Y-loops A, B, and C respectively ^8^. The loop-like structures observed for several TEs were reminiscent of Y-loops, leading us to ask whether these TE transcripts originate within Y-loop genes.

To test this idea, we used RNA FISH against TEs together with visualizing Y chromosome gene transcripts. We found that three of our five TEs exhibited clear colocalization with transcripts from Y chromosome genes. *Juan* probes illuminate a part of *kl-3* positive Y-loop B (Figure 2A) and *HMS Beagle* colocalizes with *kl-5* positive Y-loop A (Figure 2B). While *kl-2* has not previously been shown to form a Y-loop, we found that and *accord2* colocalizes with *kl-2* which creates a small loop pattern as well (Figure 2C). These observations indicate that TEs may be localized within the introns of these genes. Consistently, males lacking the Y chromosome (*XO*) showed significant reduction of loop-like signal of these three TEs (Figure 2D-I), confirming that these TE transcripts originated from the Y chromosome.

**Figure 2.**
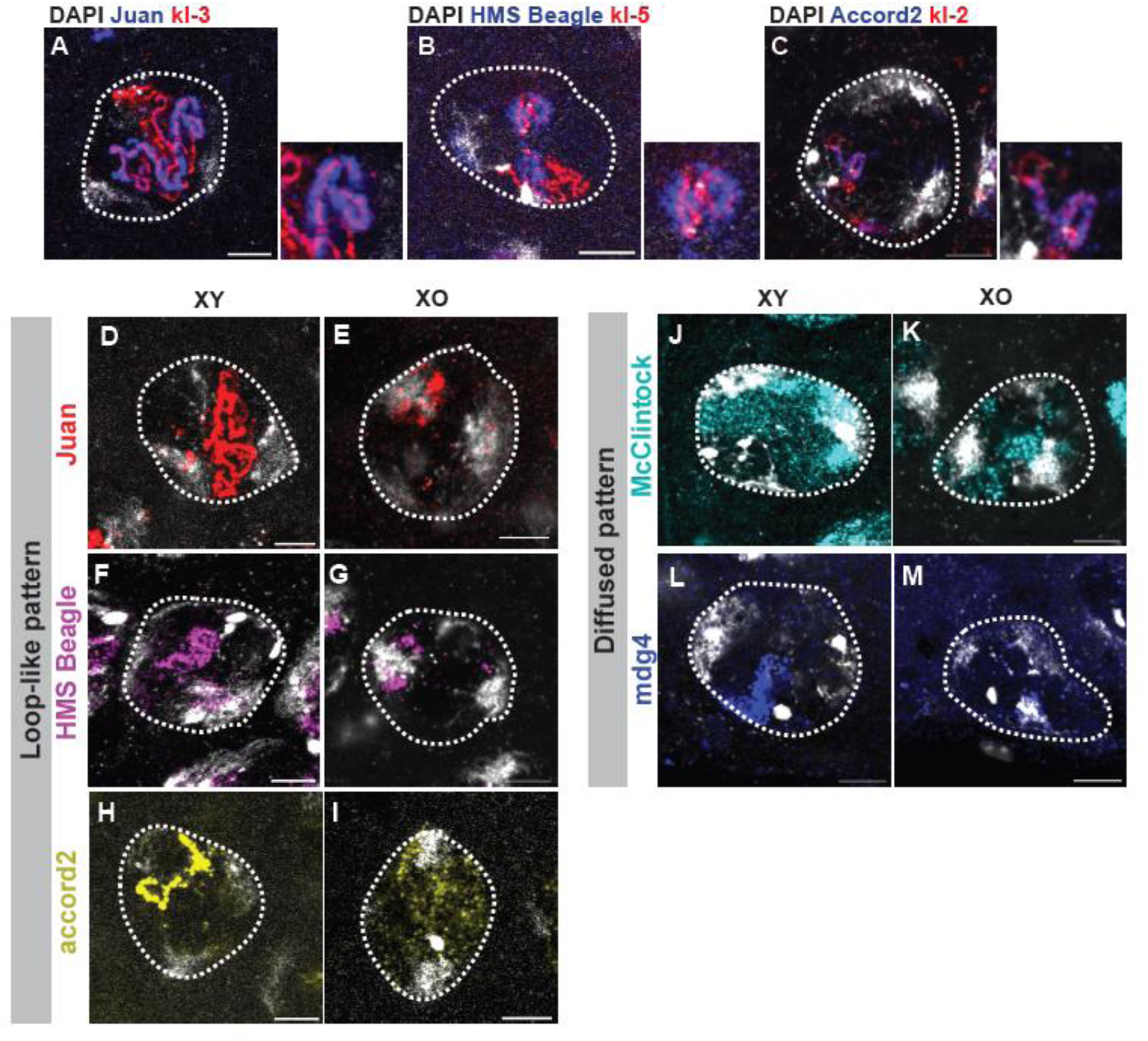
TE transcripts originate from the Y chromosome. (**A-C**) Representative RNA FISH images of late SCs (S5–S6) using indicated probes. (**D-M**) Representative RNA FISH images of late SCs (S5–S6) using indicated probes for XY males (*w1118*) or XO males. DAPI (gray). White dotted lines encircle SC nuclei. All scale bars: 5 µm.

Although we did not detect clear colocalization of the other two TEs, which exhibited a diffuse rather than loop-like localization pattern, with Y-loops, we observed a marked reduction in their diffuse RNA signals in *XO* flies (Figure 2J-M).

These findings indicate that a substantial fraction of the TE transcripts detected in spermatocytes originates from the Y chromosome.

### The majority of TEs expressed in SCs are intronic TEs

Previous studies have shown that these Y chromosome fertility genes are co-transcriptionally spliced, and splicing mutants demonstrated a significant drop in late exon transcription^14^. This drop was particularly apparent after giant introns located in these genes, indicating a difficulty in transcribing these regions^14^. We used *bamGal4* mediated knockdown of a splicing factor, *U2af38* (*bam>UAS-U2af38^HMS^*^04505^) to attenuate splicing of Y chromosome fertility genes. *U2af38* RNAi led to a significant reduction in transcripts from all five TEs (Figure 3A-K). Furthermore, RNA sequencing of isolated SCs revealed a reduction in the transcript levels of a large fraction of TEs, even though most TEs themselves are not subject to splicing (Figure 3L). It should be noted that the reduction detected by RNA sequencing (Figure 3L) is less pronounced for some TEs than that measured by RNA FISH (Figure 3K). We therefore reanalyzed the RNA-seq data using Telescope, a locus-level quantification method ^27^ (Figure S1A), and further filtered the dataset to exclude reads likely derived from degraded TE transcripts (Figure S1B), reasoning that the discrepancy might arise from the detection of partial or degraded TE RNAs by RNA sequencing. Although these analyses altered the apparent expression patterns of several TEs, they still did not fully recapitulate the reductions observed by RNA FISH. Further studies will be necessary to determine the source of this discrepancy and to develop analytical approaches that more accurately quantify TE transcripts.

**Figure 3.**
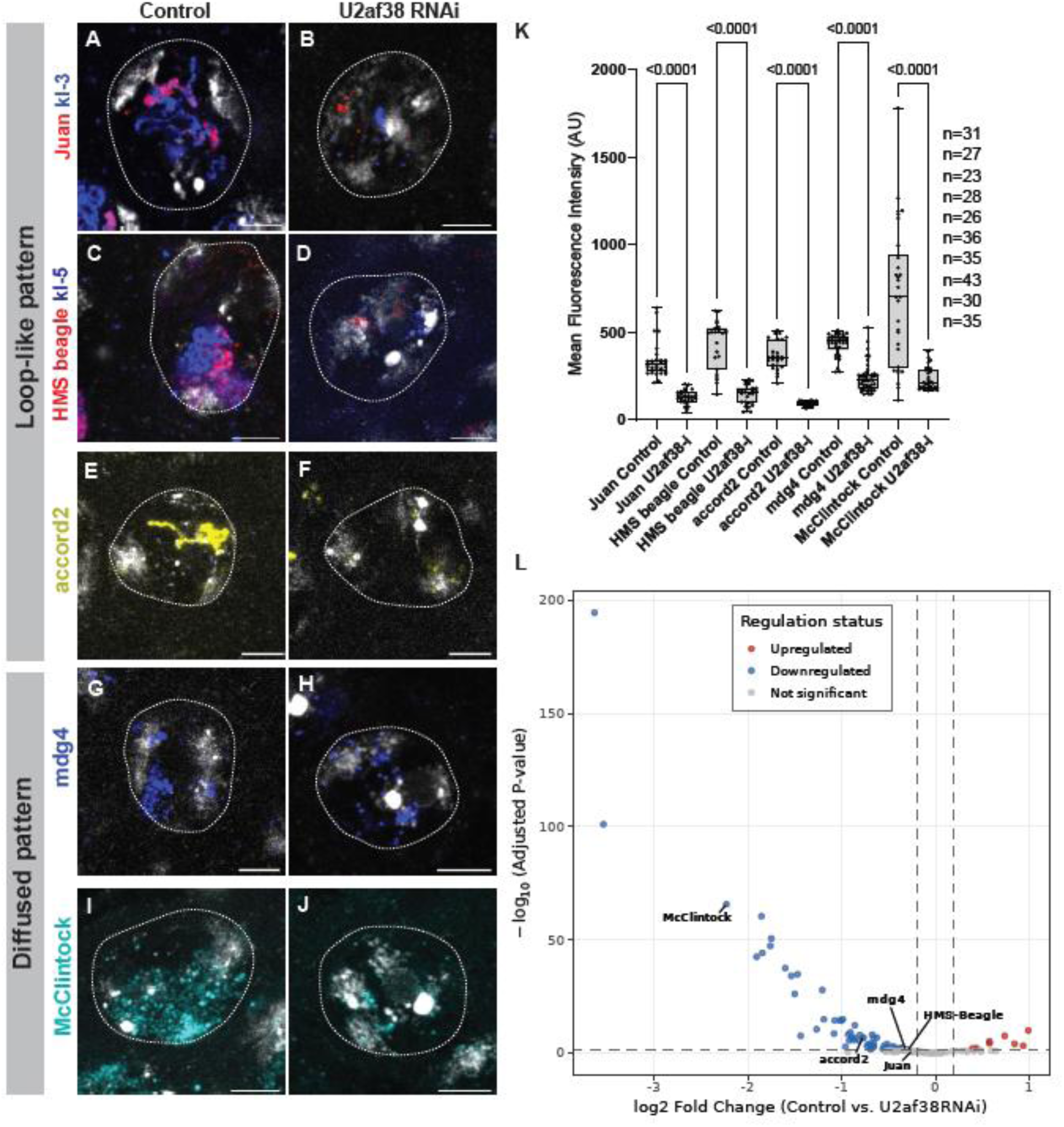
The majority of TEs expressed in SCs are intronic TEs. (**A-J**) Representative RNA FISH images of late SCs (S5–S6) hybridized with the indicated probes in testes from control (*bamGal4*) flies or *bamGal4*-driven U2af38 RNAi flies. (**K**) box plots showing the 25–75% range (box), median (center line), and minimum to maximum (whiskers), with all data points which are quantified intensities measured in the nuclear region of late SCs (S5–S6) from indicated genotypes. n indicates the number of scored nuclei. p-values were calculated using Šídák’s multiple comparisons test and are shown in the top of the graph. (**L**) Volcano plot showing DESeq2 analysis of RNA-seq data from isolated SCs comparing control (*bamGal4*, Blanks-GFP) and U2af38 knockdown (*bamGal4* >U2af38 RNAi, Blanks-GFP) testes using Kallisto pseudoalignment and quantification with TE consensus sequences. Features were colored by regulation status: upregulated (adjusted p-value < 0.05, log2 fold change > 0.2), downregulated (adjusted p-value < 0.05, log2 fold change < −0.2), or not significant. Only TEs are shown in this volcano plot. Differentially expressed genes are shown in Figure S2. Data shown in these plots are provided in Spreadsheet 3. White dotted lines encircle SC nuclei. DAPI (gray). All scale bars: 5 µm.

Despite these discrepancies in the magnitude of the reduction, both RNA FISH and RNA sequencing consistently detected decreased TE transcript levels in U2af38 RNAi samples. While there may be an indirect effect of global changes in transcription due to splicing disruption, the combination of the close association of several TE transcripts with fertility gene transcripts and the significant reduction of TE transcripts when splicing is perturbed indicates that the high expression of the majority of TEs in SCs is likely due to transcriptional readthrough of intronic insertions.

### *Juan* is expressed from *kl-3* intron 5

We identified three TEs which colocalize with specific Y chromosome genes by RNA FISH (*kl-5: HMS Beagle, kl-3: Juan,* and *kl-2: accord2*). Based on the reduction of RNA FISH signal observed in *U2af38* RNAi, we hypothesized that these TEs are illuminating nascent transcript of Y chromosome host genes. To better understand the location from which TEs are transcribed, we focused on *Juan* and attempted to map its exact location.

First, we examined the localization of Juan in the absence of *kl-3*. To accomplish this, we used a set of reciprocal translocations between the X and Y chromosomes ^28^. By crossing females which contain a distal breakpoint on the Y chromosome with males which contain a proximal breakpoint, male progeny will lack the intervening region ^29^. In flies lacking *kl-3*, *Juan* RNA FISH signal is drastically reduced (Figure 4A, B), indicating that Juan is expressed from the *kl-3* locus.

**Figure 4.**
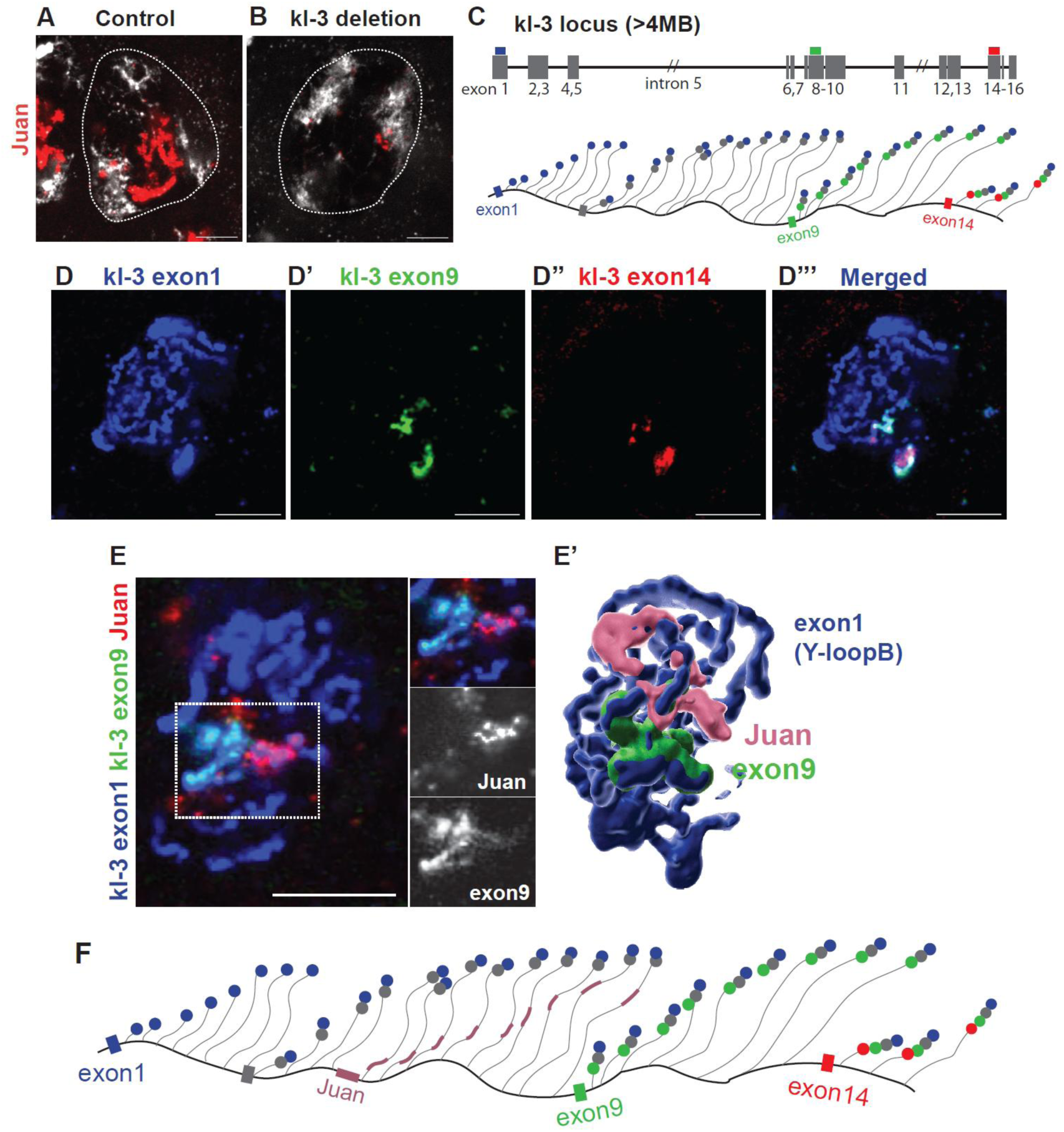
*Juan* is expressed from *kl-3* intron 5. (**A, B**) Representative RNA FISH images of late SCs (S5–S6) from indicated genotypes using a probe against *Juan*. (**C**) A schematic of the *kl-3* locus and a model illustrating the regions of nascent *kl-3* transcripts recognized by the indicated probes. (**D-D’’’**) Representative RNA FISH images of late SCs (S5–S6) using indicated probes. (**E, E’**) Representative RNA FISH images of late SCs (S5–S6) using indicated probes (**E**) and 3D rendering using Imaris software (**E’**). (**F**) A possible interpretation of observed pattern and predicted location of *Juan* insertion. White dotted lines encircle SC nuclei. DAPI (gray). *w1118* was used for all images. All scale bars, 5 µm.

Next, we examined the colocalization of *Juan* RNA with *kl-3* RNA more closely. While the *Juan* and *kl-3* exon 1 probes colocalize in certain regions, they do not share a perfect overlap. *kl-3* exon 1 signal becomes visible earlier in SC development than *Juan,* indicating that *Juan* is located later than exon 1. Even after both signals become visible, there are regions of *Juan* and *kl-3* which do not colocalize, likely representing both transcript which does not yet include the *Juan* insertion and transcript post-splicing of the *Juan* containing intron (Figure 2A). To better understand the location of the *Juan* insertion, we used additional probes against later *kl-3* exons (Figure 4C). As expected, the exon 1 probe labeled most of the *kl-3* loop, whereas the exon 9 probe labeled only a portion of the loop. The exon 14 probe labeled an even smaller region that was contained within the exon 1/exon 9 double-positive domain (Figure 4D-D’’’), consistent with the model shown in Figure 4C. Next, we mapped the position of *Juan* within the *kl-3* transcription unit using these exon probes. We found that the Juan signal did not colocalize with either the exon 9 or exon 14 probe, indicating that *Juan* is located within an intronic region between exon 1 and exon 9 (Figure 4E, E’). Because the *Juan* probe labels a substantial portion of Y-loop B, the *Juan* insertion is likely located a considerable distance upstream of the following exon. If the following exon were immediately adjacent to the *Juan* insertion, the intron would likely be spliced out soon after *Juan* transcription, and *Juan* RNA FISH signal would no longer remain tightly associated with the *kl-3* loop. Therefore, we conclude that *Juan* is most likely located in the middle of the longest intron upstream of exon 9 in the *kl-3* gene, corresponding to intron 5 (Figure 4F).

### *Juan* localization dramatically varies in closely related strains

It is important to note that we found some stocks in which TE localization patterns differ compared to *w1118*. We checked wildtype *D. melanogaster* strains for their *Juan* localization patterns and found that most strains have similar TE RNA FISH patterns to *w1118*, except for *Hikone-R*. In *Hikone-R* SCs, *Juan* diffuses throughout the Y-loop B organizing region (Figure 5 A-C). Although rare, this secondary localization pattern can be observed in random fly stocks maintained in our laboratory, including Stock #137 (*UASp-FRT-H3-GFP-PolyA-FRT-H3-mKO/3* ^30^) (Figure 5D). To better understand how this altered TE pattern arises, we isolated the Y chromosome from Stock #137 in our *w1118* stock background and found that the Y chromosome from this stock was sufficient to induce diffusion of Juan in the *w1118* background, indicating there may be changes in the Y chromosome which result in altered expression or insertions of *Juan*.

**Figure 5.**
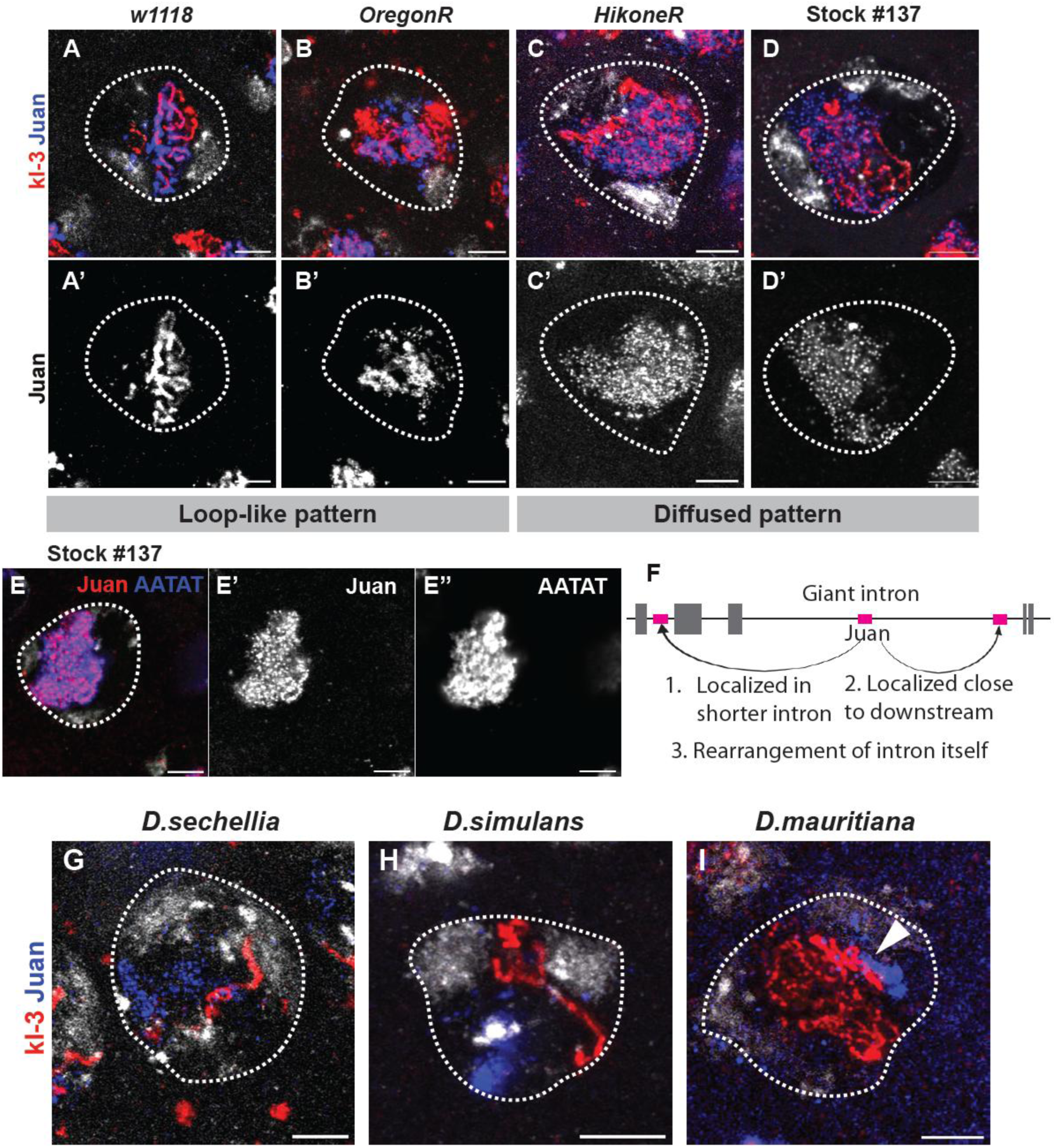
*Juan* localization dramatically varies in closely related strains. (**A-D**) Representative RNA FISH images of late SCs (S5–S6) from indicated genotypes using indicated probes. **A’-D’** panels show only Juan channel of **A-D**. (**F**) Possible interpretation of observed diffused *Juan* pattern seen in **C** and **D**. (**G-I**) Representative RNA FISH images of late SCs (S5–S6) of indicated genotypes using indicated probes. DAPI (gray). White dotted lines encircle SC nuclei. All scale bars: 5 µm.

The diffuse localization pattern of *Juan* resembles the intronic signal observed using a probe against Y-loop B intronic AATAT repeats in a previous study^14^. Therefore, we examined the colocalization of *Juan* RNA and AATAT repeat transcripts and found that they completely overlapped (Figure 5E). These findings raise the possibility that *Juan*, while still transcribed from a *kl-3* intron in these rare stocks, may be located in a different intron, or a different place within the same intron, which is retained similarly to the intronic AATAT repeat transcripts in contrast to most strains of *D. melanogaster.* (Figure 5F). Further studies will be required to determine the molecular basis of this altered localization pattern.

Given that TE localization patterns can vary even within *D. melanogaster*, we sought to examine how TE expression and localization patterns have changed in closely related non-*melanogaster* species. *D. melanogaster* and its closest relatives in the *Drosophila simulans* clade (*Drosophila sechellia*, *Drosophila simulans*, and *Drosophila mauritiana*) diverged ∼250 thousand years ago ^31,32^. Although the exons of Y chromosome fertility genes are highly conserved between these species, the giant introns differ substantially in size and repeat composition^15^. These species exhibit hybrid male sterility in reciprocal crosses^31^ and a recent study has shown that hybrid sterility is caused by fertility factor splicing defects^15^. *kl-3* RNA FISH signals in *D. sechellia* and *D. simulans* exhibited a loop-like pattern in both species similar to that observed in *D. melanogaster*. However, the loops were much less prominent and appeared shorter, likely reflecting the change in intron sizes or splicing dynamics. *Juan* displayed a patch-like localization pattern either adjacent to the chromosome or within the nucleoplasm and but never observed to colocalize with the *kl-3* loop (Figure 5G, H). *D. mauritiana* exhibited a *kl-3* loop of similar size to that observed in *D. melanogaster* (Figure 5I). Although Juan signals were closely associated with the *kl-3* loop, they did not overlap with the loop itself (Figure 5I).

The extensive variation in TE-containing fertility genes indicates that these loci are subject to strong selective pressure. This observation is consistent with the previously documented rapid evolution of Y-linked loci and suggests that TEs may contribute to the genetic diversification of these giant fertility genes.

## Discussion

In this study, we examined the stage-specific expression and localization of TE RNA in SCs. We found that TE RNA stays within the SC nucleus and adopts element-specific localization patterns. For three of the five TEs we examined, their nuclear localization represents colocalization with fertility genes which contain giant, repeat-rich introns. A previous study has reported that 412 retrotransposon transcripts localize to Y-loop A (*kl-5*), demonstrating that TE RNAs are components of the giant Y-loop transcriptional domains rather than dispersed nuclear transcripts^20^. Moreover, using single-cell RNA sequencing analysis, a recent study identified a coordinated burst of transposable element expression during primary spermatocyte differentiation that coincides with activation of the giant Y-linked fertility genes, suggesting that Y-loop transcription creates a permissive environment for TE expression^5^. However, direct evidence supporting these hypotheses as an explanation for the massive accumulation of TE transcripts in spermatocytes has been still lacking. In this study we demonstrated colocalization of specific TEs with three of the giant kl genes (*kl-5: HMS Beagle, kl-3: Juan,* and *kl-2: accord2)* in *D. melanogaster*. Further, we showed that TE RNA transcripts were reduced upon perturbation of splicing and Y chromosome gene transcription, indicating that these TEs are likely transcribed from Y introns. We used a combination of exon-specific probes for *kl-3* to map the location of the *Juan* insertion in *kl-3* and found that *Juan* is likely located in a giant intron (intron 5) of the *kl-3* gene. Finally, we examined variation in TE localization patterns within *D. melanogaster* and across closely related species, finding that TE localization is not conserved in the *D. simulans* clade and even in some strains of *D. melanogaster*.

A more complete assembly and annotation of the Y chromosome will help to understand the mechanism of TE localization and changes in closely related strains. While significant advances in assembling the *Drosophila* Y chromosome have been made recently^17,33^, large portions of the giant introns of the Y chromosome remain unknown. Using existing assemblies, we were able to locate three *HMS Beagle* insertions in *kl-5* locus, but could not locate any *Juan* insertion in *kl-3* locus (Spreadsheet 1). This could highlight flaws in previous bioinformatic analyses of the co-transcription of TEs with Y chromosome genes. Not only would short read RNA sequencing not detect fusion transcripts from TEs far into the giant introns, but DNA sequencing would indicate that the TEs are not present when the introns contain gaps of unknown sequence which may be hiding TEs. Our TE probes may present a valuable tool for examining the behavior of giant introns which cannot yet be resolved with sequencing technologies.

It’s interesting to note that *Juan* and *HMS Beagle* do not behave similarly to a previously described Y-loop intronic TE, *412*, in *XO* males ^20^. While *412* RNA, which normally colocalizes with *ORY (ks-1)*, is almost nonexistent in *XO* SCs ^20^, *Juan* and *HMS Beagle* remain present but with altered localization. Both of these TEs localize to an autosome in the absence of the Y-loops (Figure 2D-G). It has been shown previously that loss of the Y chromosome affects global chromatin states and transcription^34^. It’s possible that insertions of *Juan* and *HMS Beagle* which are not present on the Y chromosome become activated in the absence of the Y chromosome, and that this activation is prevented if parts of the Y chromosome remain, such as in deletion of *kl-3* alone (Figure 4A, B). Further work will be needed to elucidate the mechanism by which this change occurs.

We have shown that TE localization differs across certain lines of *D. melanogaster*. In most *D. melanogaster* stocks, *Juan* signal remains tightly associated with the thread-like signal from probes detecting exon 1 of *kl-3*. In certain backgrounds, including *Hikone-R* and our stock #137: *UASp-FRT-H3-GFP-PolyA-FRT-H3-mKO/3*, Juan signal is diffused around the Y-loop B region, remaining separate from loops A and C. The Y chromosome from stock#137 was sufficient to induce this altered phenotype, indicating potential structural changes to the Y chromosome which lead to this pattern.

The diffuse localization pattern of *Juan* completely overlaps with the intronic signal observed using a probe against intronic AATAT repeats in a previous study ^14^, suggesting the possibility that the location of the *Juan* insertion in these stocks may differ, causing *Juan* RNA to be processed similarly to AATAT (Figure 5F). This also suggests that spliced introns containing TE transcripts together with satellite RNA transcripts may persist in spermatocytes for extended periods. Whether these TE transcripts are translated into viral particles and remain capable of active mobilization is currently unknown. Although we did not detect cytoplasmic localization of these TE RNAs, it will be important to investigate their potential activity and determine whether they contribute to the rapid evolutionary diversification of Y-linked fertility gene loci observed among closely related *Drosophila* species in this and previous studies.

Taken together, our findings suggest that TEs contribute to the genetic diversification of giant Y-linked fertility genes. We propose that the continual turnover of repetitive elements within Y-linked introns provides a mechanism through which rapidly evolving repetitive DNA shapes the structure and regulation of these genes, with potential consequences for germline gene expression, male fertility, and ultimately, reproductive isolation and speciation.

## Methods

### Fly husbandry and strains

Flies were raised on standard Bloomington medium (Lab Express) at 25°C and young flies (0- to 7-day-old adults) were used for all experiments. The following fly stocks were obtained from the Bloomington *Drosophila* Stock Center (BDSC); *w1118* (BDSC3605), U2af38 RNAi HMS04505 (BDSC57585), *HikoneR* (BDSC4267), *OR-A* (BDSC99765), *T(1;Y)V24/C(1)A* (BDSC3060), *C(1)A;T(1;Y;3)W27* (BDSC4049), *bamGal4/X* (BDSC80579).

*kl-3* deletion males were created by crossing females from *T(1;Y)V24/C(1)A* (BDSC3060) with males from *C(1)A;T(1;Y;3)W27* (BDSC4049) (see Goldstein et al. 1982^29^ for a detailed description of the crossing scheme).

*C(1)RM/C(1;Y)/0* was a gift from George Watase. All *XO* males were created using *C(1)RM/0* females and *w1118* males.

Stock #137 *(UASp-FRT-H3-GFP-PolyA-FRT-H3-mKO/3)* was a gift from Xin Chen ^30^. *Blanks-GFP* (FBal0263998) was a gift from Yukiko Yamashita ^13^. *Drosophila simulans* w501, *Drosophila sechellia*, and *Drosophila mauritiana* were gifts from Barbara Mellone.

### RNA fluorescence in situ hybridization

#### Single molecule RNA FISH

Testes were dissected in 1X phosphate buffered saline (PBS) and then fixed in 1 ml of 4% formaldehyde/PBS for 45 minutes. Fixed testes were rinsed 2 times with 1 ml of 1X PBS, then resuspended in 1 ml of chilled 70% ethanol, and incubated for 1 hour – overnight at 4 °C. Testes were rinsed briefly in 1 ml of wash buffer (2X SSC (ThermoFisher) and 10% deionized formamide), then incubated overnight at 37 °C in the dark with 50 nM of probe in 100 µl of Hybridization Buffer containing 2X SSC, 10% dextran sulfate (MilliporeSigma), 1 μg/μl of yeast tRNA (MilliporeSigma), 2 mM vanadyl ribonucleoside complex (NEB), 0.02% RNase-free BSA (ThermoFisher), and 10% deionized formamide. Then, testes were washed 2 times for 30 minutes each at 37 °C in the dark in 1 ml of prewarmed wash buffer (2X SSC, 10% formamide) and resuspended in a drop of VECTASHIELD with 4’,6-diamidino-2-phenylindole (DAPI) (Vector Lab).

Quasar 570 labeled Stellaris probe against *Juan* and Quasar 670 labeled Stellaris probes against *HMS Beagle* and *mdg4* were designed with LGC Biosearch Technologies (www.biosearchtech.com/stellarisdesigner). CAL Fluor Red 610 labeled Stellaris probes against *accord2* and *McClintock (quasimodo2)* were a kind gift from Christopher Ellison ^5^. Quasar 670 labeled Stellaris probe against *accord2* was obtained from LGC Biosearch Technologies using probe sequences described previously ^5^. Quasar 670 labeled Stellaris probe against *kl-3* exon 1 and Quasar 570 labeled Stellaris probe against *kl-5* exons 1-6 was obtained from LGC Biosearch Technologies using probe sequences described previously^13^. The AATAT satellite transcript probe was obtained from Integrated DNA Technologies. Probe and target sequences are provided in Spreadsheet 2.

#### Hybridization chain reaction (HCR) RNA FISH

Testes were dissected in 1X PBS and then fixed in 1 ml of 4% formaldehyde/PBS for 30 minutes. Fixed testes were washed for five minutes each in 1 ml 1X PBS + 0.2% Triton X-100 (Thermo Fisher) (PBST), 1 ml 50/50 PBST/5X SSC + 0.1% Tween-20 (5XSSCT), 1ml 5XSSCT. Testes were prehybridized in 100 μl Hybridization Buffer (Molecular Instruments) at 37°C for 30 minutes. Testes were then incubated overnight at 37 °C in the dark with 20 nM of probe in 50 µl of Hybridization Buffer. Testes were washed 15 minutes in 1 ml 100% Probe Wash Buffer (Molecular Instruments), then 5 minutes each in 1ml 75% Probe Wash Buffer/25% 5XSSCT, 1ml 50% Probe Wash Buffer/50% 5XSSCT, and 25% Probe Wash Buffer/75% 5XSSCT, all at 37°C. Testes were washed 5 minutes with 1ml 5XSSCT, then 5 minutes with 200 μl Amplification Buffer (Molecular Instruments) at room temperature. During the wash steps, hairpins were prepared by heating in a PCR machine at 95°C for 90 seconds, then cooling at room temperature in the dark for 30 minutes. Testes were then incubated at room temperature 6 hours – overnight in 50 μl Amplification Buffer with 6 pmol hairpins. After incubation, testes were washed four times with 1ml 5XSSCT, two 5 minute washes, one 30 minute wash, and one 5 minute wash, before resuspension in a drop of VECTASHIELD with DAPI.

HCR (v3.0) probes against *HMS Beagle* and *kl-2* exons 1-2 were designed by Molecular Instruments. Target sequence information can be found in Spreadsheet 2.

To combine HCR FISH with smFISH, the HCR FISH protocol was used. smFISH probes were added during the probe hybridization step at the same concentration used for smFISH.

Images were taken using a Zeiss LSM800 confocal microscope with airyscan module with 63X oil immersion objective (NA = 1.4) using Zen software. Images were processed by image J/FIJI.

### Quantification of TE transcript

Image-J/Fiji was used for image quantification. Z-stack series were taken using a 63X/1.4 NA oil objective on a Zeiss LSM800 confocal microscope with airyscan module. Images were Airyscan processed with standard strength. Average RNA FISH intensity was measured in SC nuclei using a single slice from a z-stack, and background level within the same testis was subtracted. The same acquisition setting was used across all samples.

### SC sorting and RNA-sequencing

*Blanks-GFP* flies were raised at 25°C. Testes were dissected in 1X Phosphate buffered saline (PBS) in a glass spot plate. In the spot plate, each testis was carefully separated from accessory organs and placed into a 1.7 mL microcentrifuge tube containing 1mL of 1% BSA in PBS supplied with RNaseOut (Invitrogen) and kept on ice.

After collection of 30∼40 testes/tube, the supernatant was removed and the pellet of testes was rinsed with 1mL of PBS two times. 100 µL of dissociation buffer (1X PBS with 2mg/mL collagenase (Sigma), 4 mg/mL dispase II (Millipore), 0.5 mM MgCl_2_, 0.5 mM CaCl_2_) was added. Then, samples were incubated at 37 °C for 15 minutes; the tubes were tapped every 5 minutes to aid in dissociation. Tubes were put on ice immediately after dissociation. 400 µL of 1% BSA/ PBS was added to the tube, and the sample was pipetted up and down using a 29G insulin syringe to dissociate the cells. The dissociated cells were then pipetted through a 125 µm metal mesh into tubes for Fluorescence-activated Cell Sorting (FACS). Samples were kept on ice before proceeding to cell sorting.

Cells were sorted using the BD FACSymphony S6 SE with a large diameter (130 µm) nozzle and using 4-way precision purity mode. Cells collected from wild-type testes were used as a negative control. GFP-positive cell populations were gated according to the negative control. Sorted cells were collected in a 1.7 mL microcentrifuge tube filled with 0.3 mL of collection buffer. RNA was extracted using RNA Extraction Protocol – RNeasy Plus Micro Kit (Qiagen). Ribosomal RNA (rRNA) depletion, library preparation, and 40M paired-end sequencing on the Element AVITI were performed by AmpSeq (https://www.ampseq.com/). RNA sequencing data is deposited in NCBI GEO under accession number GSE337949.

### RNA-sequencing data analysis

#### Read Quality Control and Preprocessing

We assessed raw paired-end RNA-seq reads for quality using FastQC v0.12.1. Adapter sequences and low-quality bases were trimmed using TrimGalore v0.6.7 with a quality threshold of Q20 and a minimum read length of 36 bp.

#### Gene Expression Quantification

Trimmed reads were aligned to the *Drosophila melanogaster* reference genome (release 6.65) using STAR v2.7.11b. To ensure unique gene assignments, STAR was run in two-pass mode with a maximum of 1 multi-mapping location per read (--outFilterMultimapNmax 1). STAR indices were generated using the genome FASTA and gene annotation GTF (FlyBase r6.65). Using the The --outSAMstrandField intronMotif parameter, alignments were output as coordinate-sorted BAM files with annotated strand information.

Gene-level read counts were quantified using featureCounts (Subread v2.0.3) in paired-end mode with reverse strandedness (-s 2) to match the library preparation protocol. Read pairs were assigned to genes based on overlap with exonic features defined in the FlyBase r6.65 GTF annotation.

#### Transposable Element Expression Quantification

##### TE Annotation

A comprehensive transposable element annotation was generated using RepeatMasker v4.1.5 with the D. melanogaster TE consensus sequences from the Bergman laboratory curated library (https://github.com/bergmanlab/drosophila-transposons). RepeatMasker was run in sensitive mode (-s) against the D. melanogaster r6.65 genome to identify all genomic TE insertions. The RepeatMasker output was converted to GTF format, preserving both locus-level coordinates and family annotations for each TE insertion.

##### TE-Specific Alignment and Quantification

Due to the repetitive nature of TEs, we employed a separate alignment strategy to more accurately quantify TE expression. First, trimmed reads were aligned to the D. melanogaster r6.65 genome using STAR v2.7.11b with parameters optimized for multi-mapping reads: --outFilterMultimapNmax 150 and --winAnchorMultimapNmax 150. Since many TE insertions share highly similar nucleotide patterns, allowing multi-mapping is necessary to avoid losing the vast majority of reads. These alignments were output as unsorted BAM files, and subsequently name-sorted using samtools to ensure compatibility with downstream TE quantification tools.

To maintain locus-level resolution, TE expression was quantified using Telescope v.1.0.3 ^27^ which employs a Bayesian framework to reassign multi-mapped reads to their most likely TE of origin based on local alignment context. Telescope was run with the following parameters: --attribute gene_id for locus-level quantification, --reassign_mode conf for confidence-based probabilistic reassignment, --conf_prob 0.95 to retain only alignments with ≥95% posterior probability, --overlap_threshold 0.2 requiring reads to overlap at least 20% of their length with a TE annotation. We chose this approach over simpler counting methods (e.g. featureCounts, HTSeq) because Telescope explicitly models the uncertainty in multi-mapped read assignments, reducing false positives from ambiguous alignments while preserving TE-mapping reads that would otherwise be discarded. Since Telescope outputs locus-level counts for individual TE insertion, we aggregated these counts by TE family using the family classifications from the RepeatMasker-derived GTF (summation of counts sharing classification).

##### Kallisto Pseudoalignment and Quantification

As an additional methodological control, we repeated alignment (in this case, pseudoalignment) and quantification of transcripts using kallisto/0.50.1.To enable parallel quantification of gene transcripts and transposable element families within a single “competitive” framework, we constructed a combined pseudo-alignment reference by concatenating the Drosophila melanogaster FlyBase transcript sequences (dmel-all-transcript-r6.65.fasta) with the same 127 TE consensus sequences as aforementioned. A kallisto index was built from the combined reference using kallisto index. Transcript-level abundance estimation was performed for each sample using kallisto quant in paired-end mode with 100 bootstrap resampling iterations (-b 100). Gene-level count estimates were subsequently aggregated from transcript-level output using tximport ^35^ with a transcript-to-gene mapping derived from the FlyBase r6.65 GTF. TE-level estimated counts were extracted directly from the abundance.tsv files by filtering for entries corresponding to the 127 TE consensus sequences, merged into a single count matrix across all four samples, and rounded to integers before DESeq2 was run.

##### Normalization

As a note: standard DESeq2 normalization (median-of-ratios ^36^) relies on the assumption that most of the features across the transcriptome are not differentially expressed and that fold changes are symmetric. As discussed above, U2af38 (U2 snRNP Auxiliary Factor) is an essential, universally conserved splicing factor required for recognition of the polypyrimidine tract at the 3’ splice site and for stable recruitment of the U2 snRNP during the first step of pre-mRNA splicing. Thus, global depletion of U2af38 via RNAi is expected to broadly impair pre-mRNA splicing, producing widespread intron retention, misrecognition of alternative splice sites, and activation of nonsense-mediated decay (NMD) for aberrantly processed transcripts. Such transcriptome-wide perturbations may violate the assumption underlying median-of-ratios normalization that most genes exhibit no systematic change in abundance, potentially biasing size-factor estimation. Applying default DESeq2 normalization under these conditions could cause the algorithm to artificially rescale RNAi sample counts upward, generating false-positive “upregulated” features and potentially masking the true magnitude of transcriptome-wide mRNA depletion.

To circumvent this bias, we normalized all count matrices against reads derived from pre-ribosomal RNA (pre-rRNA) transcribed spacer sequences. Ribosomal RNA genes are transcribed by RNA Polymerase I (Pol I) from the nucleolar organizer rDNA locus. Entirely independent of the U2AF-dependent RNA Polymerase II splicing pathway, the Pol I primary transcript (the “45S” pre-rRNA) is processed co- and post-transcriptionally into the mature 18S, 5.8S, 2S, and 28S ribosomal RNA species. This processing removes and rapidly degrades three spacer elements embedded in the primary transcript: the 5’ External Transcribed Spacer (ETS) and the two Internal Transcribed Spacers (ITS1 and ITS2). Because these spacers are intrinsically short-lived and do not accumulate to the levels of the stable mature rRNAs, their steady-state abundance in a total RNA-seq library reflects the active Pol I transcription rate rather than the ribosome-resident RNA pool ^37^. Pol I transcription and pre-rRNA processing are mechanistically independent of U2af38 and the spliceosome, making ETS/ITS reads an appropriate internal reference anchor for normalization in the context of a global splicing factor knockdown.

##### Pre-rRNA Spacer Read Quantification

Reads from each sample were aligned to the Drosophila melanogaster ribosomal DNA reference sequence (GenBank: M21017.1 ^38^) using Bowtie2 ^39^ in end-to-end sensitive mode, with unaligned reads suppressed (--no-unal --end-to-end --sensitive). The M21017.1 reference was treated as a standalone alignment target entirely separate from the BDGP6/dm6 genome index used for standard transcript quantification, ensuring rDNA-mapping reads were quantified without competition from the primary genome alignment. Resulting alignments were coordinate-sorted and indexed using SAMtools (Li et al., 2009).

Spacer-region read counts were extracted from each sample’s BAM file using samtools view restricted to three non-overlapping genomic intervals defined from the M21017.1 GenBank feature annotation: the 5’ External Transcribed Spacer (ETS; positions 10,866–11,726), Internal Transcribed Spacer 1 (ITS1; positions 1,996–2,721), and Internal Transcribed Spacer 2 (ITS2; positions 2,903–3,287). Secondary alignments and unmapped reads were excluded via the combined bitflag filter -F 260. Counts were restricted exclusively to these three spacer intervals; the mature 18S (1–1,995), 5.8S (2,722–2,844), 2S (2,873–2,902), and 28S (3,288–7,232) rRNA coding sequences were deliberately excluded, as these regions encode stable, highly abundant, ribosome-incorporated RNA species whose steady-state levels do not reflect active transcription rate and would introduce rRNA-pool-dependent bias into the normalization.

Combined ETS/ITS1/ITS2 read counts per sample were as follows: BlanksGFP_sorted1, 421,785; BlanksGFP_sorted2, 577,682; U2af38 RNAi_sorted1, 1,653,947; U2af38 RNAi_sorted2, 1,982,050. Spacer-mapping reads constituted 3.79% and 4.98% of total reads in the BlanksGFP replicates, compared to 14.90% and 13.95% in U2af38 RNAi replicates—a consistent approximately 3–4-fold elevation that we confirmed to be independent of sequencing depth (BlanksGFP total read pairs: 11,136,966 and 11,600,949; U2afRNAi total read pairs: 11,098,528 and 14,209,451). This systematic elevation in pre-rRNA spacer abundance in knockdown spermatocytes likely reflects either compensatory upregulation of Pol I transcriptional output in response to global splicing stress, or impaired pre-rRNA processing with consequent accumulation of unprocessed spacer-containing intermediates, and is discussed further in the Results.

##### Size Factor Calculation and DESeq2 Model Fitting

Custom DESeq2 size factors were derived directly from the combined ETS/ITS1/ITS2 spacer read counts. For each sample, the raw spacer count was divided by the geometric mean of all four samples’ spacer counts, yielding per-sample size factors: BlanksGFP_sorted1, 0.446; BlanksGFP_sorted2, 0.611; U2afRNAi_sorted1, 1.750; U2afRNAi_sorted2, 2.097. These factors were assigned directly to the DESeqDataSet object via the sizeFactors() slot prior to model fitting, bypassing DESeq2’s internal estimateSizeFactors() step. Because DESeq2 detects pre-populated size factors and skips its own estimation procedure, all subsequent dispersion estimation and hypothesis testing used exclusively the pre-rRNA-derived scaling.

We analyzed differential expressions in parallel DESeq2 models, each using the same pre-rRNA size factors but differing in the quantification methods mentioned above. First, gene-level counts produced by featureCounts ^40^ from STAR-aligned reads were supplied directly to DESeq2 via DESeqDataSetFromMatrix(). Second, transcript-level abundances from kallisto ^41^ were aggregated to gene level using tximport ^35^ with a transcript-to-gene mapping derived from the FlyBase GTF; the resulting gene-level count estimates were rounded to integers and supplied to DESeq2 via DESeqDataSetFromMatrix() with the same pre-rRNA size factors. The TE Kallisto abundance estimates were extracted, rounded, and supplied to DESeq2 as well. Next, transposable element (TE) family-level counts from Telescope ^27^ were analyzed in a DESeq2 model using unfiltered family counts. And finally, an independent TE analysis was performed on divergence-filtered Telescope counts as described below. All models used the design formula ∼condition, contrasting U2af38 RNAi against BlanksGFP controls. Fractional count estimates from Telescope and kallisto were rounded to the nearest integer prior to DESeq2 input. Differential expression was tested using the Wald test; p-values were corrected for multiple comparisons using the Benjamini–Hochberg false discovery rate procedure^42^. We considered features with an adjusted p-value < 0.05 and absolute log2 fold change > 0.2 statistically significant.

##### Divergence-Based Transposable Element Filtering

To focus TE expression analysis on younger, potentially active insertions and reduce signal contribution from ancient, likely transcriptionally inert copies, we subjected Telescope locus-level counts to a post-hoc divergence-based filter prior to a parallel family-level DESeq2 analysis. Per-locus percent divergence from the TE consensus sequence was retrieved from the divergence attribute field of the Telescope TE annotation GTF, which records the Kimura 2-parameter divergence estimated by RepeatMasker for each annotated TE insertion. Loci whose divergence from consensus exceeded 5% were excluded from downstream quantification. We selected this threshold to retain insertions with high sequence similarity to their respective consensus sequences, which are more likely to represent recently transposed, transcriptionally competent copies, while excluding diverged remnants whose expression may reflect readthrough transcription or mapping artifacts rather than bona fide TE activity.

Reads assigned by Telescope to exclude reads originating high-divergence loci were discarded entirely and not redistributed to other loci. This post-hoc approach confers the benefit of preventing the introduction of artifactual count reassignments that could confound family-level quantification. Following locus-level filtering, retained read counts were reaggregated to the TE family level by summing across all qualifying loci belonging to each family. The resulting filtered family-level count matrix was supplied to a separate DESeq2 model using the same pre-rRNA size factors applied to all other analyses. Gene-level counts were identical between the unfiltered and divergence-filtered analyses, as the filter operates exclusively on TE locus assignments.

##### Limitation of Methods

Please note that due to their repetitive nature and propensity to insert within long expanses of low-complexity DNA regions, many transposable elements are notoriously difficult to evaluate using bioinformatic methods. These low-complexity regions (oftentimes the Y-chromosome of *Drosophila*) are the most difficult to sequence, leading to large gaps within the genome. Downstream, this substantially limits the creation/classification of TE’s as consensus sequences, identification of insertions, and accurate alignment of transcripts. We employed multiple quantification methods, along with best practices in bioinformatics, in an attempt to mitigate these effects.

#### Visualization

Volcano plots were generated in R using ggplot2, with point labels applied using ggrepel to minimize label overlap. Features were colored by regulation status: upregulated (adjusted p-value < 0.05, log2 fold change > 0.2), downregulated (adjusted p-value < 0.05, log2 fold change < −0.2), or not significant. For gene-level volcano plots, the ten features with the highest combined significance score (−log10(adjusted p-value) × |log2 fold change|) were labeled, capturing features that are both highly significant and of biologically relevant effect size.

### Identification of TE insertions

We identified TE insertions within *kl-3* and *kl-5* by extracting each gene’s intronic intervals from the FlyBase r6.67 annotation (dmel-all-r6.67.gff) and intersecting them against the FlyBase TE annotation GTF, keeping transposable-element features fully contained within an intron. For each insertion we recorded its Y-chromosome coordinates, TE family/class, host intron, and fractional position within that intron (0 = 5’ splice site, 1 = 3’ splice site). Identified TE list is provided in Spreadsheet 1.

### Statistical analysis and graphing

No statistical methods were used to predetermine sample size. The experiments were not randomized. The investigators were not blinded to allocation during experiments and outcome assessment. All experiments were independently repeated at least three times to confirm the results. Statistical analysis and graphing were performed using GraphPad Prism 10 software. The Data are presented as box plots showing the 25–75% range (box), median (center line) and minimum to maximum (whiskers). p-values were calculated using Šídák’s multiple comparisons test. Individual numerical values displayed in a graph are provided in Source data1.

## Supporting information

Sourcedata1

## Acknowledgements

We thank Xin Chen, Christopher Ellison, Barbara Mellone, George Watase, Yukiko Yamashita, and the Bloomington Drosophila Stock Center for reagents; Evan Jellison and Li Zhu for flow cytometry support. This research is supported by R35GM128678 from the National Institute for General Medical Sciences and a start-up fund from UConn Health (to M.I.).

## Author Contributions Statement

E.K.B. and M.I. conceived the project, designed and executed experiments, analyzed data, and drafted manuscript. J.G. performed the bioinformatic analyses and wrote the Methods section. A.R. provided support for the bioinformatic analyses used for transposon identification and quantification. All authors edited the manuscript.

---

**Figure S1.**
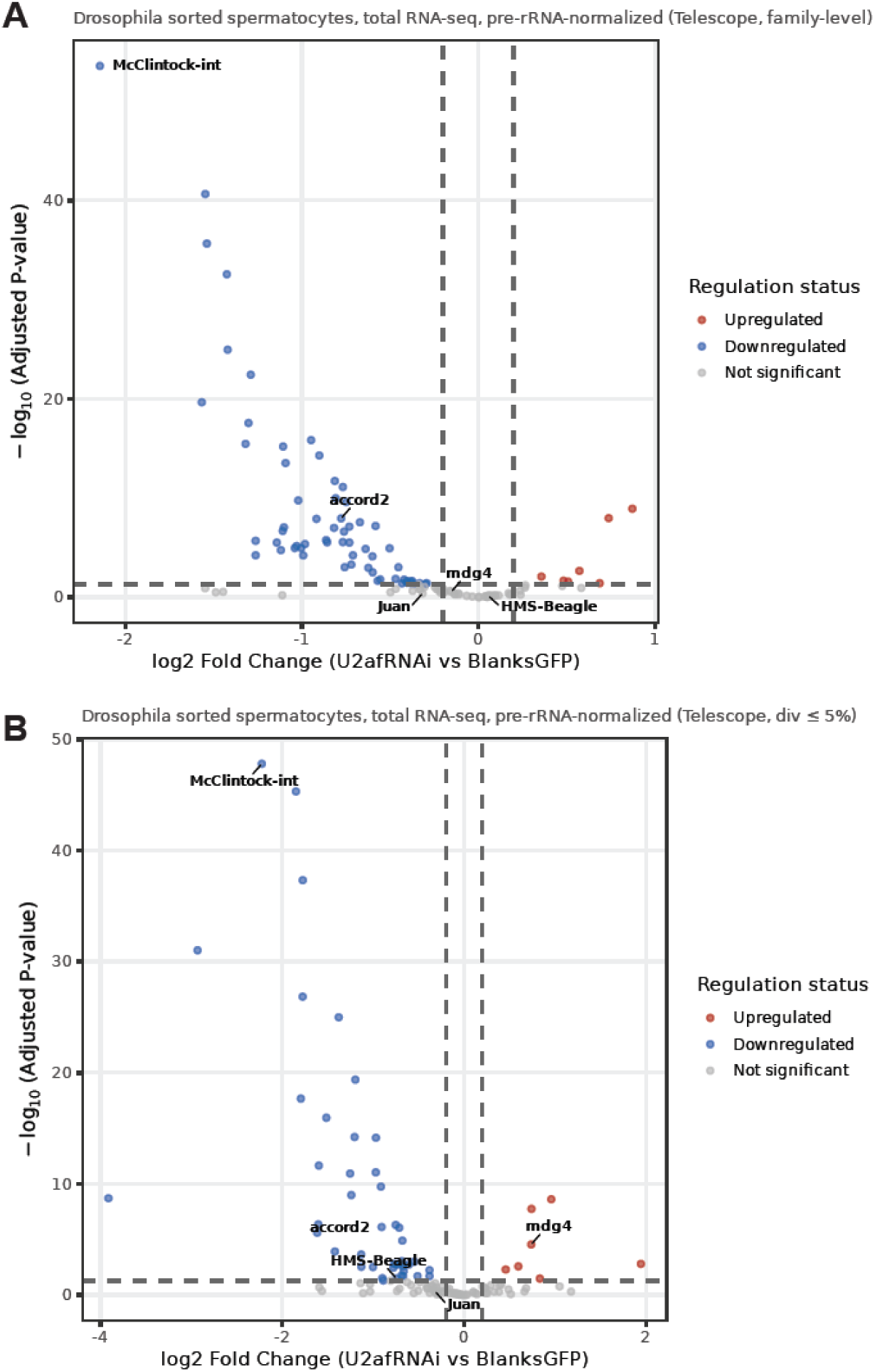
Telescope based quantification of TE expression. (**A-B**) Volcano plot showing DESeq2 analysis of RNA-seq data from isolated SCs comparing control (*bamGal4*, Blanks-GFP) and U2af38 knockdown (*bamGal4* >U2af38 RNAi, Blanks-GFP) testes using Telescope. Features were colored by regulation status: upregulated (adjusted p-value < 0.05, log2 fold change > 0.2), downregulated (adjusted p-value < 0.05, log2 fold change < −0.2), or not significant. Data shown in these plots are provided in Spreadsheet 3. In B, TE aligned reads whose divergence from consensus exceeded 5% were excluded.

**Figure S2.**
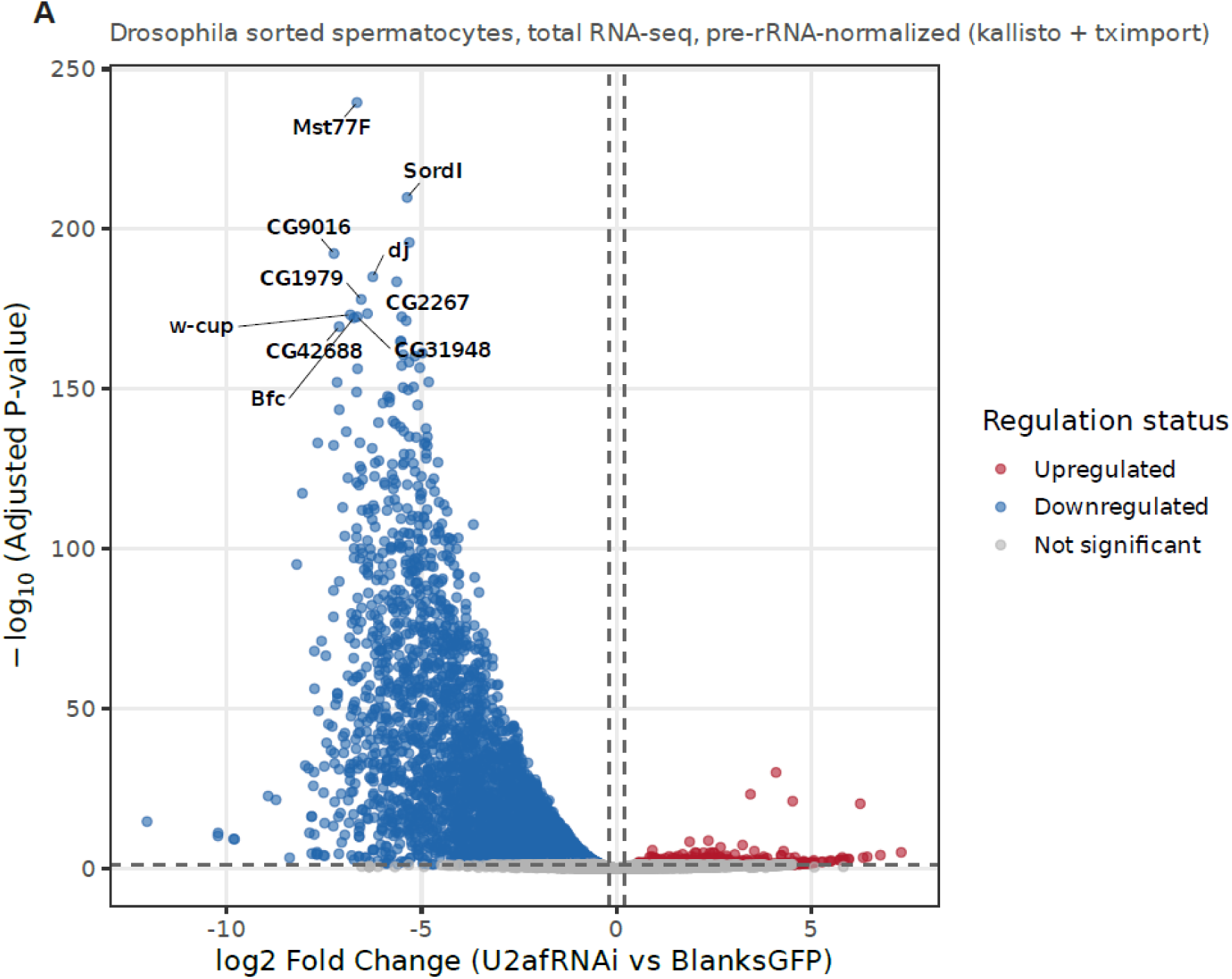
Gene expression changes in SCs from U2af38 knockdown testes. Related to Figure 3L. (**A**) Volcano plot showing DESeq2 analysis of RNA-seq data from isolated SCs comparing control (*bamGal4*, Blanks-GFP) and U2af38 knockdown (*bamGal4* >U2af38 RNAi, Blanks-GFP) testes using featureCounts. Features were colored by regulation status: upregulated (adjusted p-value < 0.05, log2 fold change > 0.2), downregulated (adjusted p-value < 0.05, log2 fold change < −0.2), or not significant. Only genes are shown in this volcano plot. Data shown in these plots are provided in Spreadsheet 3.

